# Quantitative imaging of gene therapy delivery vehicles using CEST-NMR/MRI

**DOI:** 10.1101/2022.11.09.515733

**Authors:** Bonnie Lam, Mark Velasquez, A.J. Velasquez-Mao, Kevin Godines, Wissam AlGhuraibawi, Michael Wendland, Jeff Pelton, Moriel Vandsburger

**Author notes:** Corresponding Author: Moriel Vandsburger, PhD, 284 Hearst Memorial Mining Building, Department of Bioengineering, University of California, Berkeley, CA, 94721.

## Abstract

**Purpose:** Gene therapy employing AAV vector-mediated gene delivery has undergone substantial growth in recent years with promising results in both preclinical and clinical studies, as well as emerging regulatory approval. However, the lack of methods for quantifying the efficacy of gene therapy from cellular delivery of gene editing technology to specific functional outcomes remains an obstacle for the efficient development of gene therapy treatments. Building upon prior works that utilized a genetically encoded Lysine Rich Protein as a chemical exchange saturation transfer (CEST) reporter, we hypothesized that AAV viral capsids may generate endogenous CEST contrast from the large number of surface lysine residues.

**Methods:** Water-suppressed NMR and NMR-CEST experiments were performed on isolated solutions of AAV serotypes 1-9 on a Bruker 800MHz vertical scanner. A series of in vitro experiments were performed for thorough testing of NMR-CEST contrast of AAV2 capsids under varying pH, density, biological transduction stage, and later across multiple serotypes and mixed biological media. Reverse transcriptase (RT)-polymerase chain reaction (PCR) was used to quantify virus concentration. Subsequent experiments determined the pH-dependent exchange rate and optimized CEST saturation schemes for AAV contrast detection at 7 T.

**Results:** NMR-CEST experiments revealed CEST contrast up to 52% for AAV2 viral capsids between 0.6-0.8 ppm. Evaluation of CEST contrast generated by AAV2 demonstrates high levels of CEST contrast across a variety of chemical environments, concentrations, and saturation schemes. AAV2 CEST contrast displayed significant positive correlations with capsid density (R^2^>0.99, P<0.001), pH (R^2^=0.97, P=0.01), and viral titer per cell count (R^2^=0.92, P<0.001). Transition to a preclinical field strength yielded up to 11.8% CEST contrast following optimization of saturation parameters.

**Conclusion:** AAV2 viral capsids exhibit strong capacity as an endogenous CEST contrast agent and can potentially be used for monitoring and evaluation of AAV vector-mediated gene therapy protocols.

## 1. Introduction

Over the last decade the field of gene therapy has rapidly expanded to include in vivo somatic cell gene editing^1^. Increasingly, adeno-associated virus (AAV) vectors have been exploited as delivery vehicles for gene editing machinery such as CRISPR thanks to non-pathogenicity, organ specificity, replication defectiveness, and low immunogenicity^2^. Clinical and preclinical studies utilizing AAV vectors have explored the efficacy of different serotypes for targeting a wide range of disease models across multiple organs, including Duchenne Muscular Dystrophy (DMD)^3–5^, hemophilia^6–8^, and Alzheimer’s Disease (AD)^9,10^. Notably, gene therapy drugs utilizing AAV vectors have been gaining traction for commercial use in patients. Namely, Glybera^11^ was approved in Europe in 2012 for treatment of lipoprotein lipase (LPL) deficiency; whereas Luxturna^12^ and Zolgensma^13^ were FDA-approved in 2017 for treatment of Leber congenital amaurosis (LCA) and in 2019 for treatment of spinal muscular atrophy (SMA), respectively. Additional studies that have been conducted for the design of novel viral vector variants with enhanced organ tropism and for optimization of packing, delivery, and antibody evasion^14–16^ have collectively further enhanced the potential of AAV vectors for gene therapy.

While methods for delivering gene editing machinery and more precisely editing target mutations have vastly improved, the reliance upon invasive biopsies for verification of successful gene editing and longitudinal monitoring of spatiotemporal expression patterns remains a boundary to successful translation. Dependence upon biopsies increases the likelihood of spatial sampling error, inflicts pain, and may result in complications including sudden death when performed in the heart^17,18^. In practice, there remains a significant gap in measurement during crucial intermediate phases of gene editing. Current standards for evaluation of success or failure of a particular gene therapy protocol primarily consist of administration of a specified dose of an AAV vector followed by assessment of organ system level functional outcomes. This input-output approach towards monitoring gene therapy treatments fails to track the multiple stages of biological events involved in AAV vector-mediated gene editing including initial transduction, endosomal transport, endosomal escape, and subsequent transgene expression^19^ that take place prior to observable differences in functional outcomes. Further, in the absence of a system level functional outcome, the inability to quantify the aforementioned biological stages limits the ability to understand the point of therapeutic failure. Thus, the ability to measure delivery, editing, and transgene expression is a high profile need in gene editing.

In prior studies, we and others have demonstrated the ability to track transgene expression of the Lysine Rich Protein (LRP), a genetically engineered chemical exchange saturation transfer (CEST)-MRI reporter peptide^20^. Based on the exchange of saturated magnetization at 3.76ppm between amide protons in Lysine and surrounding bulk water, substantial CEST contrast was generated by LRP following AAV9 mediated delivery to the mouse myocardium. A potential method for monitoring the cellular delivery of AAV capsids containing gene editing machinery is to exploit similarities between AAV capsid surface composition and LRP^2021^ ^21^. Specifically, compared to the 50 lysine residues that comprise the LRP^20^, each AAV2 capsid contains over 1,000 lysine residues on its capsid surface^22,23^ that we hypothesized may generate similar contrast on CEST-MRI. In this study, we first probed whether AAV2 capsid proteins generate endogenous CEST contrast analogous to LRP through a series of in vitro experiments. The parameter space was fully explored using NMR CEST experiments under varying pH, density, biological transduction stage in cellular systems, and later across multiple commercially available AAV serotypes and mixed biological media. Subsequent experiments determined the pH-dependent exchange rate and optimized CEST saturation schemes for AAV contrast detection at 7T.

## 2. Methods

### 2.1. NMR Spectroscopy at 800 MHz

#### 2.1.1. Conventional NMR imaging of AAV2

Conventional NMR experiments were performed on AAV2 (5.23×10^8^ viral genomes/μL) in solution using an 800 MHz vertical bore system with a TXI probe (Avance I; Bruker, Ettlingen, Germany). The sample was maintained at pH 7.0 and 37°C. Additional water-suppressed NMR experiments were conducted under the same conditions. Commercially available AAV2 was purchased from Vector Biolabs and diluted in preparation for NMR experiments.

#### 2.1.2. NMR-CEST of AAV capsids in solution

In order to thoroughly test whether AAV capsids generate endogenous CEST contrast a series of experiments were first performed on preparations of AAV2 at varying concentrations and pH. First, NMR-CEST experiments were performed on AAV2 (5.26×10^8^ viral genomes/μL) in solution and a separate 0.1% poly-L-lysine (PLL) sample as a control at pH 7.0 and 37°C. Complete Z-spectra were acquired from −5 ppm to +5 ppm with a step size of 0.2 ppm across a range of saturation powers from 3.0 μT to 9.0 μT, serially acquired for all studies. CEST data were extracted as traces at the H2O resonance from a series of 1D NMR spectra collected at different saturation frequencies. Data were recorded using pulse sequence syszg2df2gppr provided by the manufacturer (Bruker BioSpin Inc., Billerica, MA.) The recycle and saturation delays were set to 17 sec and 3 sec, respectively. After saturation, the water resonance was excited using a 18° flip angle pulse, that was typically 2.5 μs in duration. The saturation and excitation portions of the sequence were separated by a Z-axis gradient pulse of 1 ms duration and at a field strength of 5 G/cm. Each scan was recorded with a total of 32 K points, using a spectral width of 12820 Hz, and with the carrier frequency set to the water resonance. Data from two scans were signal averaged for each frequency. Next, NMR-CEST experiments were performed on AAV2 samples in solution titrated to pH 6 while varying capsid densities between 5.26×10^3^ vg/μL and 5.26×10^8^ vg/μL. Imaging was performed for AAV2 samples at a concentration of 5.26×10^8^ vg/μL while varying pH between 4-7.5 (4, 5, 6, 7.5). NMR-CEST experiments were next performed on multiple AAV serotypes (1, 5, 6, 7, 9) at a concentration of 5.26×10^8^ vg/μL and pH titrated to a value of 6. All AAV were commercially available (Vector Biolabs).

#### 2.1.3 NMR-CEST of cellular transduction of AAV capsids

To test whether AAV2 generates CEST contrast during endosomal transport, HEK293T cells were transduced with AAV2 (MOI 10,000 for 2 million cells) via one hour exposure to viral media at 4°C, after which viral media was removed and replaced with regular media. Cells were harvested at 0-, 30-, 60-, 90-, and 120-minutes following removal of viral media. Control HEK293T cells were exposed to normal media for one hour at 4°C. Afterwards, endosomes were lysed, isolated, and buffered in solution at pH 6 for NMR-CEST experiments. This experiment was performed twice with identical timepoints.

#### 2.1.3. Exchange rate quantification

Samples of AAV2 at a concentration of 5.26×10^8^ vg/μL were titrated to pH values around 6 and 7 prior to imaging at 37° C. NMR-CEST Z-spectra were acquired at 111 frequency offsets between −10 ppm and +10 ppm at varying saturation powers (1, 1.5, 2, 2.5, and 3 μT). CEST data were extracted as traces at the H2O resonance from a series of 1D NMR spectra collected at different saturation frequencies. Data were recorded using pulse sequence syszg2df2gppr provided by the manufacturer (Bruker BioSpin Inc., Billerica, MA.) The recycle and saturation delays were set to 1 sec and 12 sec, respectively. After saturation, the water resonance was excited using a 18° flip angle pulse, that was typically 2.5 μs in duration. The saturation and excitation portions of the sequence were separated by a Z-axis gradient pulse of 1 ms duration and at a field strength of 5 G/cm. Each scan was recorded with a total of 32 K points, using a spectral width of 12820 Hz, and with the carrier frequency set to the water resonance. Data from two scans were signal averaged for each frequency. Z-spectral acquisitions were normalized by reference values prior multi-B_1_ fitting with Bloch-McConnell equations ^24^ to determine exchange rates for each sample.

### 2.2. Optimization at 7 T

Based on the findings of exchangeable protons from NMR-CEST experiments, we next performed CEST imaging experiments at a field strength of 7T in order to determine whether endogenous CEST contrast of AAV particles could be detected at pre-clinical relevant field strengths. Phantoms containing AAV2 (5.26×10^8^ vg/μL) were imaged at 7 T (PharmaScan; Bruker, Ettlingen, Germany) using a 2×2 surface array coil (Bruker, Billerica, MA). Complete Z-spectra acquired from −10 ppm to +10 ppm following varying values for saturation B_1_ (1, 1.5, 2, 3, 4 and 5 μT), saturation pulse duration (54.8, 36.5, 27.4, 18.3, 13.7, 11 ms), and saturation duty cycle (0.6, 0.7, 0.8, 0.85) across different combinations. A single Gaussian pulse was applied prior acquisition with a centric-ordered GRE readout sequence (TR/TE = 6.44/3.13 ms, matrix = 192 x 192, FOV = 25 mm x 25 mm, slice thickness = 3 mm, number of averages = 2). Reference acquisitions were performed before and after the acquisition of each z-spectrum and used for normalization.

In order to probe the impact of dilution in background tissue protein content, additional experiments were carried out on a phantom containing cell lysate (6.8mg protein) and AAV2 (5.26×10^8^ vg/μL). Complete Z-spectra were acquired from −10 ppm to +10 ppm following varying values for saturation B_1_ (3, 4 and 5 μT), saturation pulse duration (54.8, 36.5, 27.4 ms), and saturation duty cycle (0.7, 0.85) across all possible combinations. Image properties were identical to those previously stated.

### 2.3. CEST contrast analysis

All imaging data were analyzed using custom code written in MATLAB (MathWorks, Natick, MA). For NMR-CEST experiments, CEST contrast was calculated as MTR_asym_=[S(+ω) − S(− ω)]/S_0_).

For CEST experiments conducted at 7 T, regions of interest were drawn on reference images prior to calculation of average signal intensities for further analysis. Z-spectra were calculated by normalizing CEST acquisitions to linearly interpolated values of the −10 ppm and +10 ppm acquisitions, followed by temperature correction via linear scaling. In phantoms containing AAV in solution only, CEST contrast was quantified by 2-pool Lorentzian fitting^25^ with pools allocated for water and AAV. In phantoms containing AAV and cell lysate in solution, CEST contrast was quantified by 6-pool Lorentzian fitting with pools allocated for water, magnetization transfer (MT), amine, amide, nuclear Overhauser enhancement (NOE), and AAV. Contributions from different pools to the Z-spectrum were quantified based on a sum of Lorentzian functions as follows:

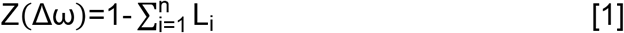

with

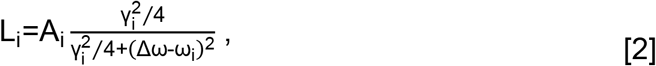

where A_i_, γ_i_, and ω_i_ represent amplitude, FWHM, and resonance frequency relative to water, respectively. Lorentzian peak amplitude values were used as the measure of CEST contrast.

### 2.4. Statistical Analysis

Regression analysis for determining correlation and significance was performed in Excel. Representative values are listed as mean ± standard deviation.

## 3. Results

Conventional (Figure 1A) and water suppressed (Figure 1B) NMR spectroscopy performed on solution of AAV2 viral capsids (5.26×10^8^ vg/μL) revealed three potential pools of exchangeable protons with adequate concentrations at frequency offsets of 3.6 ppm, 3.0 ppm, and 0.6 ppm (Figure 1B). CEST-NMR experiments were then conducted to investigate contrast levels at the aforementioned peak frequencies of AAV2 viral capsids. Z-spectra generated by AAV2 (5.26×10^8^ vg/μL) reveal broad and significant saturation transfer occurring at upfield frequencies in comparison to a solution of 0.01% PLL as shown in Figure 2. Subsequent MTRasym calculations display a strong peak around a frequency offset of 0.6 ppm. High levels of CEST contrast (~31 −52%) are maintained across the range of saturation powers tested (Supplemental Figure 1), with peak contrast increasing with saturation power prior plateauing at higher powers. On average, AAV2 generated CEST contrast of 45.6 ± 5.85% whereas PLL generated CEST contrast of 0.60±0.75%. Although goodness of fit was affected by lack of further increases in contrast with saturation power (R^2^=0.22), the relationship between CEST contrast and saturation power was still significant (P=0.02).

**Figure 1.**
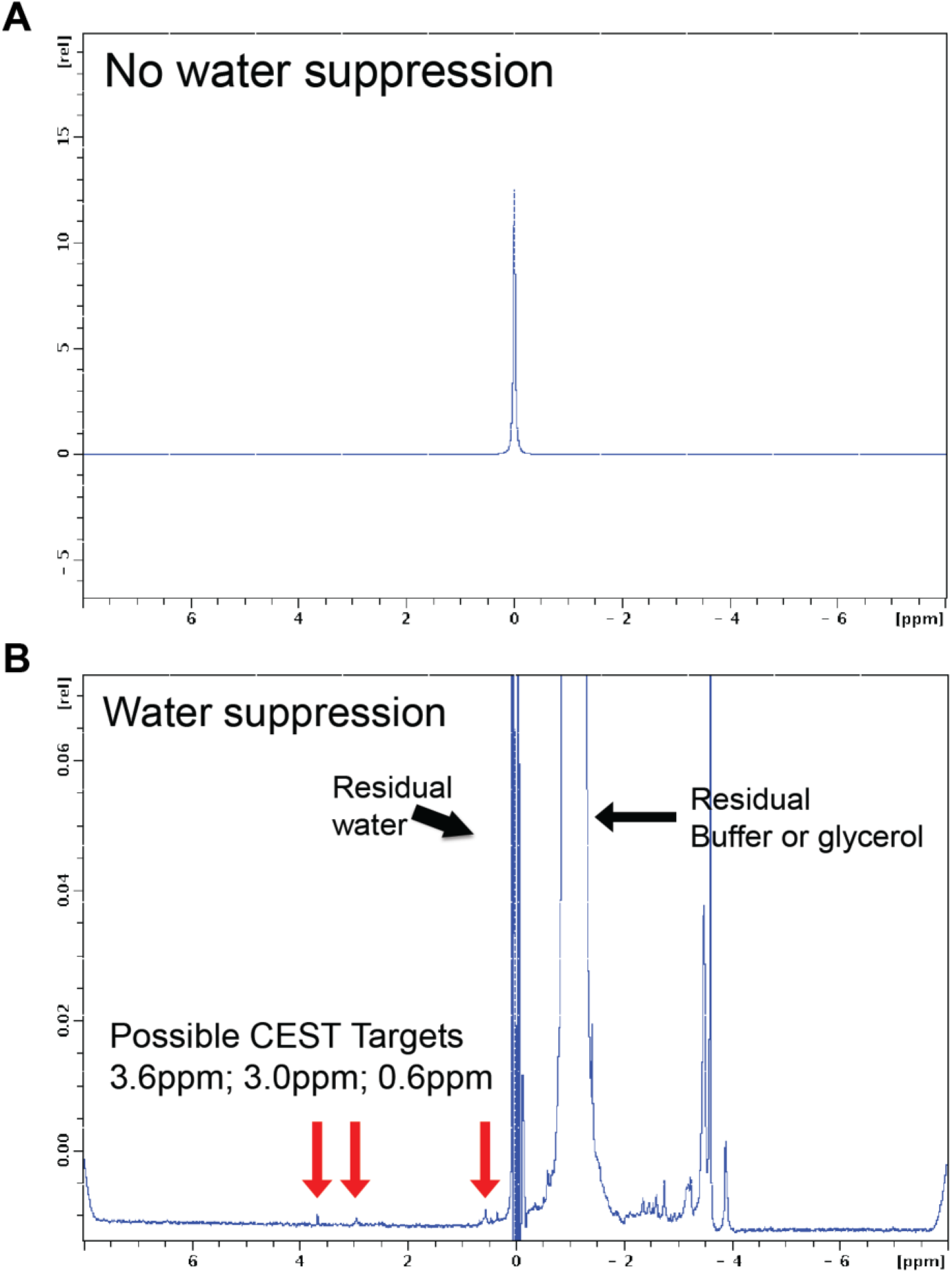
NMR Spectroscopy of AAV2 Viral Capsids. (A) Conventional NMR of AAV2 viral capsids. (B) Water-suppressed NMR of AAV2 viral capsids reveal 3 potential CEST targets at 3.6 ppm, 3.0 ppm, and 0.6 ppm.

**Figure 2.**
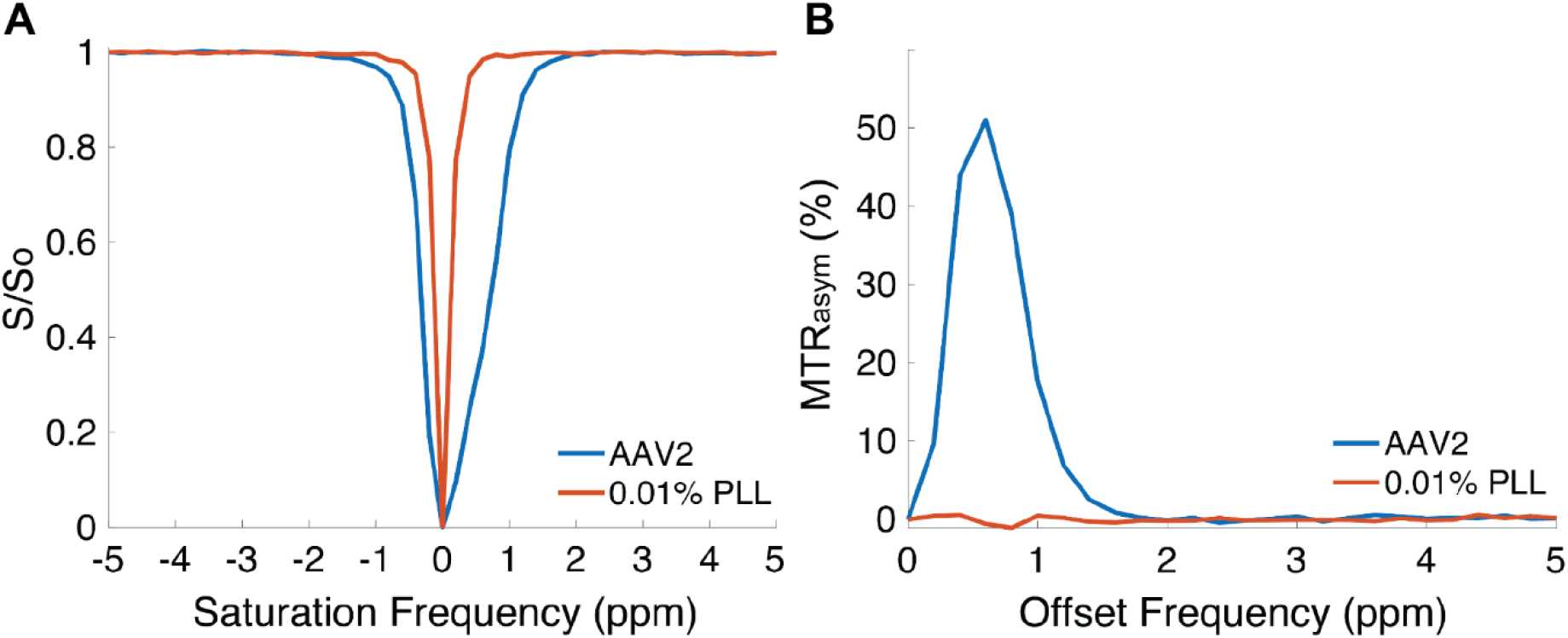
CEST-NMR of AAV2 Viral Capsids. (A) Z-spectra of AAV2 (5.26×10^8^ vg/μL) and 0.01% poly-L-lysine as a control, acquired at B_1_=6.6 μT. (B) Corresponding MTRasym analysis displays high CEST contrast for AAV2 viral capsids around 0.6 ppm.

CEST contrast of AAV was explored under different conditions of concentration, pH, and serotype. Mean CEST values for AAV2 samples across saturation powers were 0.81±0.51%, 1.04±0.84%, 3.69±1.48%, and 31.2±4.28% for concentrations of 5.26×10^3^ vg/μL, 5.26×10^5^ vg/μL, 5.26×10^7^ vg/μL, and 5.26×10^8^ vg/μL, respectively. Representative CEST contrast for varying viral capsid densities at selected low, medium, and high saturation powers are shown in Figure 3A. A strong positive correlation was demonstrated between concentration and resulting CEST contrast (R^2^>0.99, P<0.001). Differences in pH also had an effect on subsequent CEST contrast; mean CEST values for AAV2 (5.26×10^8^ vg/μL) across saturation powers were 26.3±10.4%, 31.3±11.2%, 34.5±11.1%, and 45.6±5.9% for pH values of 4, 5, 6, and 7.5, respectively. Representative CEST contrast for varying pH at selected low, medium, and high saturation powers are shown in Figure 3B. pH and CEST contrast were significantly correlated; as pH increased, CEST values also increased in most cases (R^2^=0.97, P=0.01). Evaluation of CEST contrast for different AAV serotypes yielded similar strong peaks at 0.6 ppm (Figure 4). Average CEST values across saturation powers were 33.8±5.5%, 33.4±5.7%, 28.7±4.8%, 32.4±5.9%, and 30.7±4.9% for AAV1, AAV5, AAV6, AAV8, and AAV9 respectively. Although contrast was slightly lower than the 34.5±11.1% contrast demonstrated for AAV2, robust CEST contrast was still observed across other serotypes. Interestingly, additional weaker peaks at other offset frequencies were detected for some serotypes in a power-dependent manner (Figure 4).

**Figure 3.**
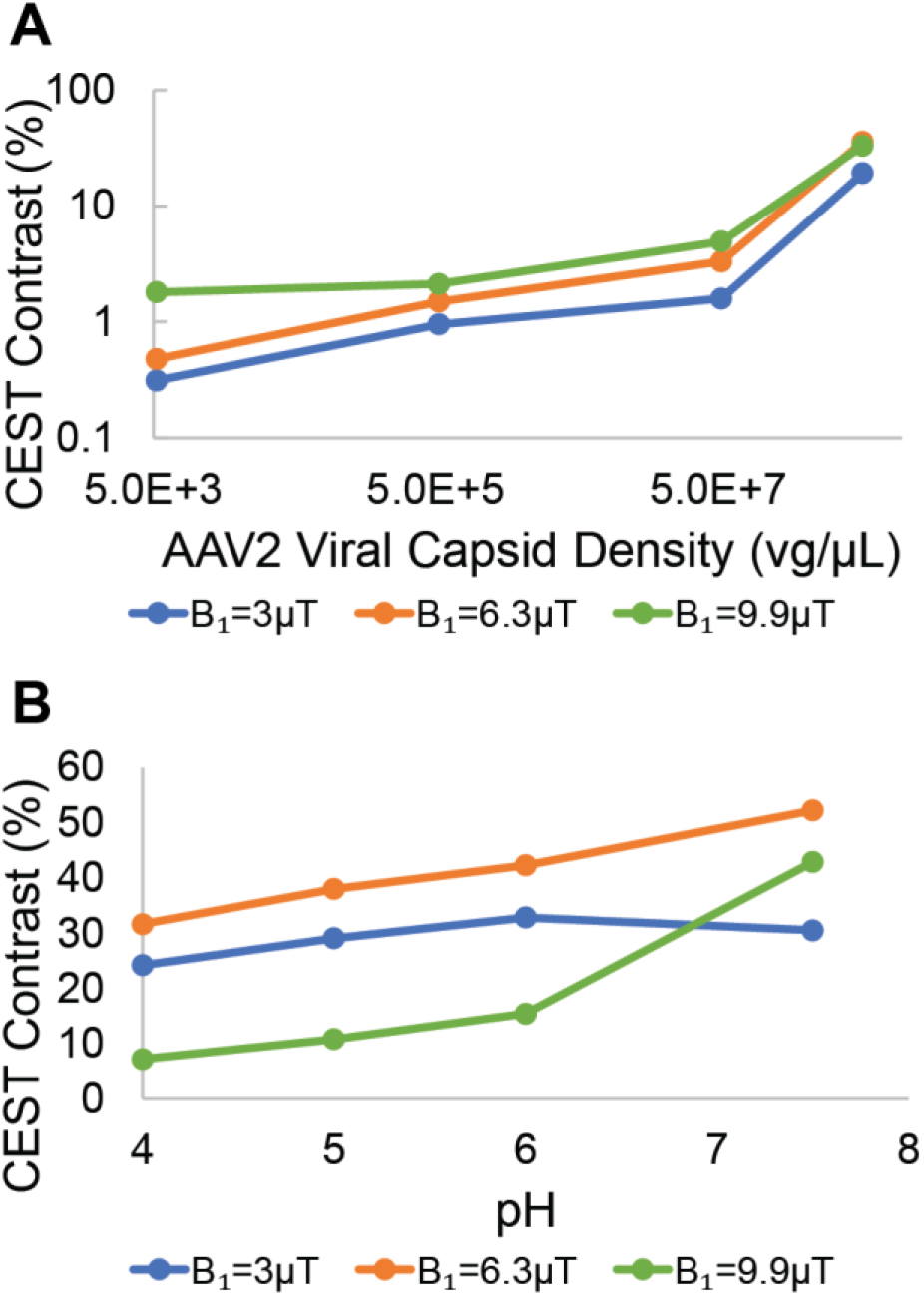
CEST Contrast of AAV2 Across Concentrations, pH, and Saturation Power. (A) CEST contrast of AAV2 viral capsids at multiple B_1_ powers while varying capsid densities from 5.26×10^3^ vg/μL to 5.26×10^8^ vg/μL. A significant increase in contrast with increasing capsid density is observed across saturation powers. (B) CEST contrast of AAV2 viral capsids (5.26×10^8^ vg/μL) at multiple B_1_ powers while varying pH values from 4 to 7.5. A general increase in CEST contrast is observed with increasing pH in most cases.

**Figure 4.**
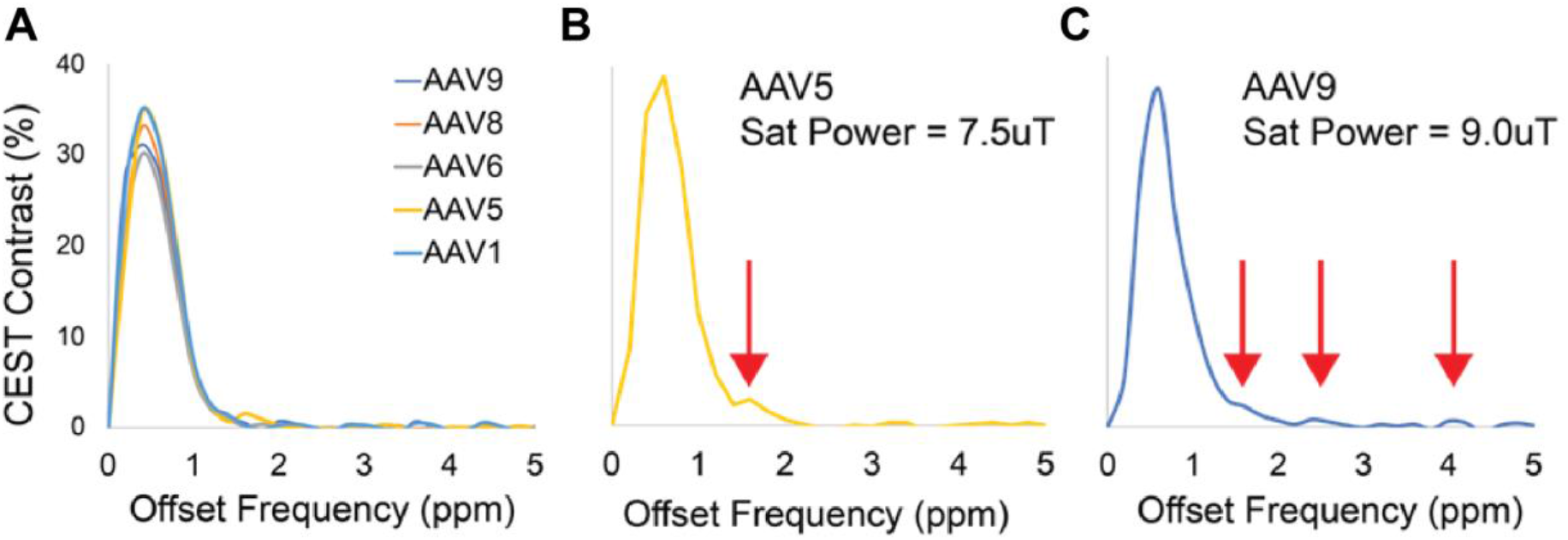
CEST Contrast Across AAV Serotypes. Repeated CEST-NMR experiments demonstrated robust CEST contrast around 0.6ppm across multiple AAV serotypes (A). Additionally, some AAV serotypes exhibited CEST contrast peaks at other offset frequencies (B,C).

Detection of stages of AAV2 internalization displayed differences in contrast across timepoints. Quantification of viral titer per cell count in the first run showed a peak value of 0.08 at T=0 minutes prior fluctuations that trended downwards to 0.03 at T=120 minutes (Figure 5A). The second run of the same experiment displayed a much different trajectory across time, with viral titer/cell count showing minor fluctuations between 0.008 at T=0 minutes and 0.01 at T=90 minutes preceding a large jump to 0.17 at T=120 minutes (Figure 5B). Viral titer/cell count for controls in both runs were near zero (<10^-6^). Corresponding measurements of CEST contrast demonstrated similar trends across timepoints, with contrast fluctuating downwards between 4.52% and 3.66% for the first run (Figure 5C) and contrast varying between 3.50% and 2.84% for the first 4 timepoints prior a jump to 6.10% at T=120 minutes (Figure 5D). Regression analysis of combined data from both runs, including controls, revealed a strong positive correlation between CEST contrast and viral titer/cell count (R^2^=0.92, P<0.001) (Figure 5E).

**Figure 5.**
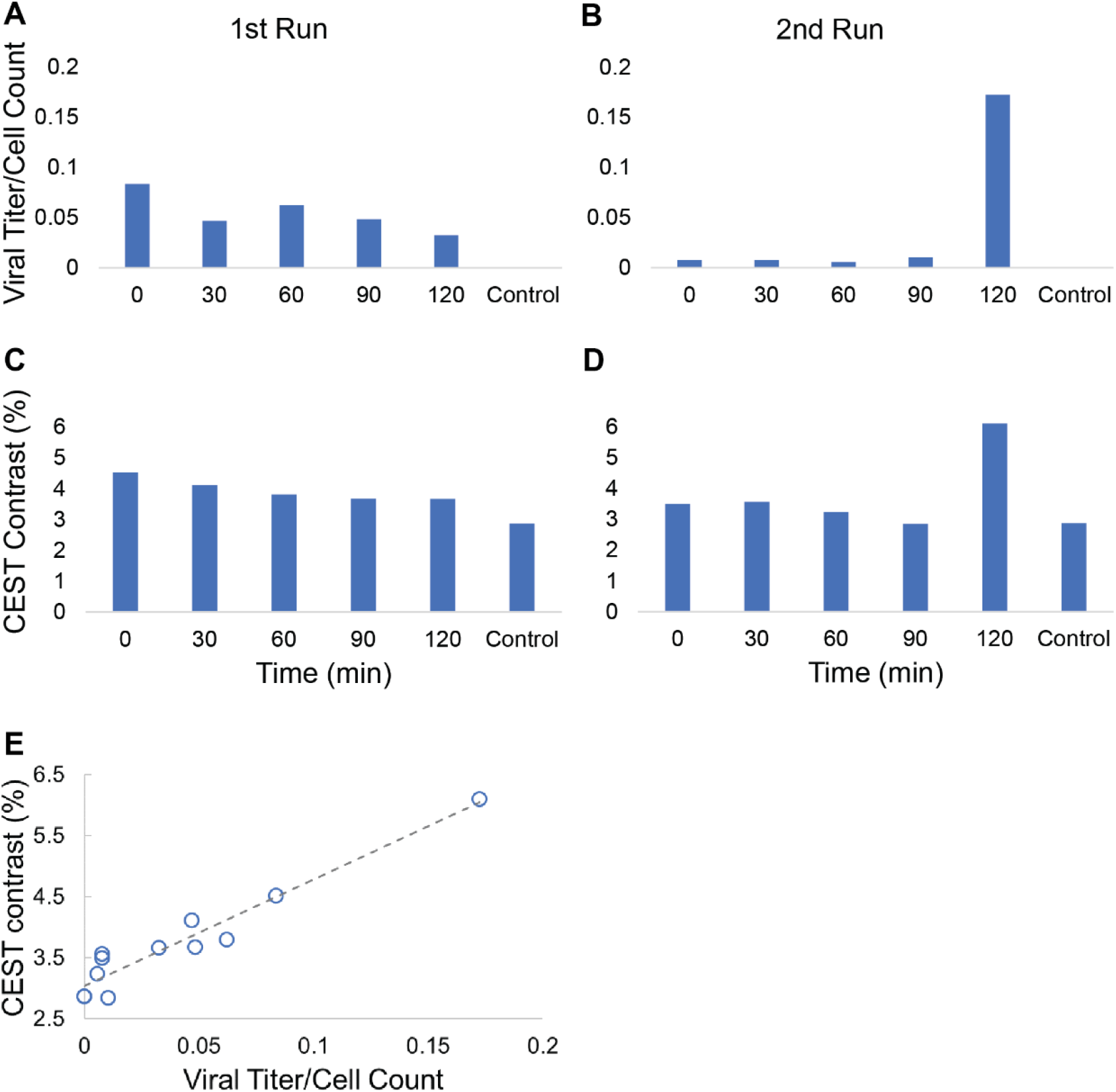
Detection of AAV2 Internalization. Repeated CEST-NMR experiments conducted on isolated endosomes harvested from transduced HEK293T cells at multiple timepoints demonstrated varying trends in viral titer per cell count and CEST contrast at 0.8 ppm. (A) Viral titer per cell count in the first run demonstrated fluctuations that gradually decreased over timepoints, whereas (B) viral titer per cell count in the second run displayed a sudden jump at 120 min. (C,D) Resultant CEST contrast in the first and second runs yielded trends that mirrored those shown in viral titer per cell count. (E) Regression analysis of combined data from both runs (including controls) revealed a strong positive correlation for CEST contrast against viral titer/cell count (R^2^=0.92, P<0.001).

Determination of exchange rates for AAV2 at varying pH yielded a mean exchange rate of 1709±433 Hz at pH values of approximately 6 and 7. More specifically, resulting exchange rates following Bloch-McConnell equation fitting were 1402 Hz and 2015 Hz for pH values of 6 and 7, respectively. Exchange rates for AAV2 demonstrated a corresponding increase with an increase in pH. Although the positive correlation between exchange rates and pH was not significant, the trend displayed is similar to the positive correlation previously observed in NMR-CEST contrast across pH values. An example of multi-B_1_ z-spectra and corresponding fitting is shown in Supplemental Figure 2.

Choices in saturation scheme parameters resulted in large differences in contrast at 7 T as shown in Figure 6. Saturation at an offset of 10 ppm represents a region far from water resonance, whereas saturation at an offset of 0 ppm represents a direct saturation of water. Similarly, a saturation offset of 0.8 ppm represents CEST contrast generation by hydroxyl protons on the AAV capsid surface. Images of an AAV2 phantom (5.26×10^8^ vg/μL) at multiple frequency offsets display corresponding differences in levels of saturation when the selected saturation scheme generates little to no contrast (Figure 6A) or high levels of contrast (Figure 6B). Initial attempts to image the AAV2 phantom used varying values of B_1_ (1-2 μT), bandwidth (150-250 Hz, corresponding to pulse durations of 11.0-18.3 ms), and duty cycle (60-80%) that resulted in very low CEST contrasts of on average 0.02±0.02%. Representative Z-spectrum and Lorentzian difference plots (Figure 6C,E) show minimal contrast generation from AAV2 capsids. Following several saturation scheme adjustments, acquisitions performed while varying B_1_ (3-5 μT), bandwidth (50-100 Hz, corresponding to pulse durations of 27.4-54.8 ms), and duty cycle (70-85%) resulted in much higher CEST contrast values that averaged 6.81±4.14%. Multiple sets of saturation schemes were found to generate very high levels of CEST contrast above 9% (Table 1). Using this optimization of saturation parameters, CEST contrast reached up to 11.8% (Figure 6D,F).

**Figure 6.**
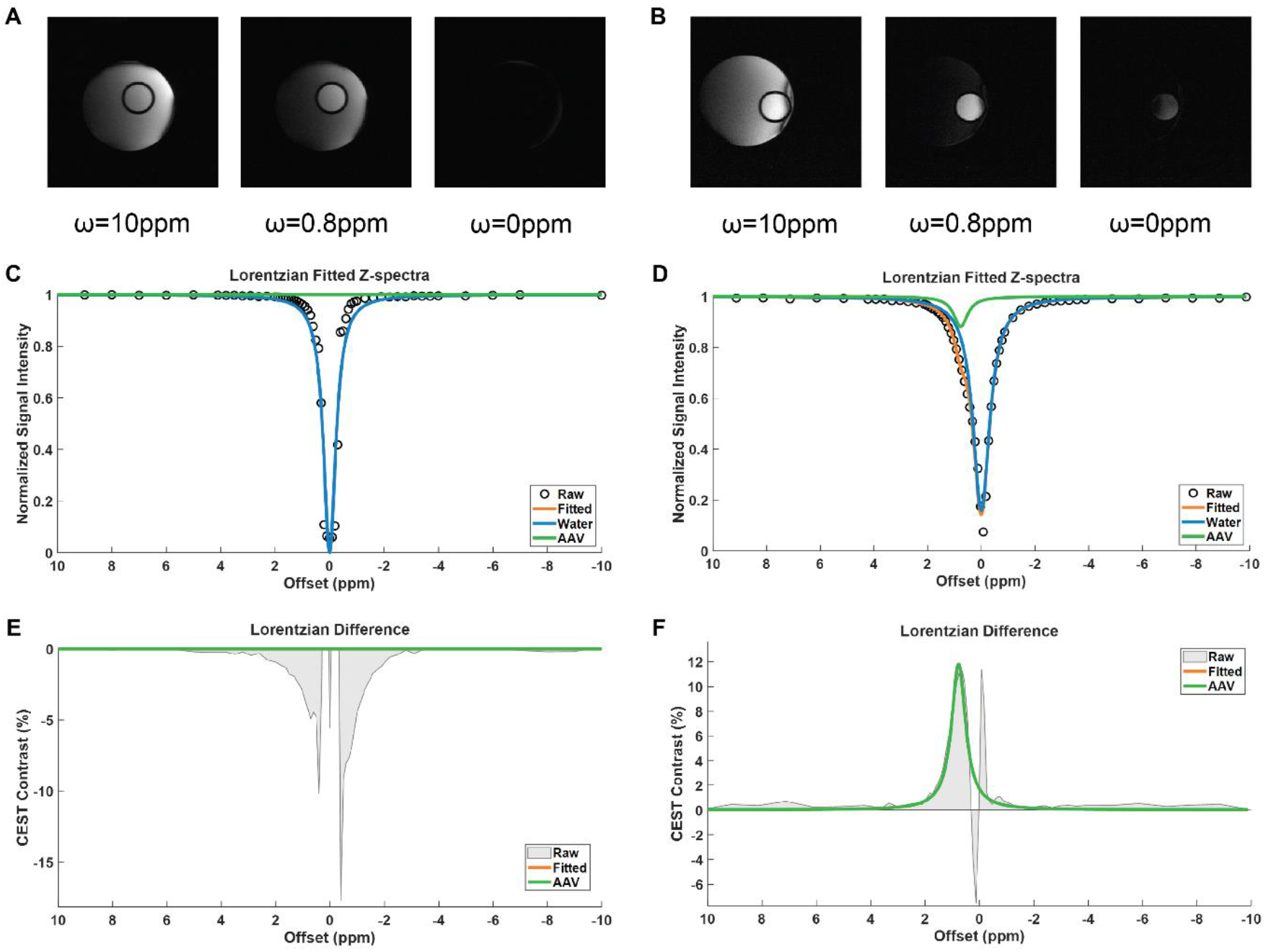
Representative Z-spectra and MTRasym of AAV2 at 7 T. (A) Images of AAV2 phantom (5.26×10^8^ vg/μL) at multiple frequency offsets that correspond to the highest acquired offset (10 ppm) and the resonant frequencies of water (0 ppm) and AAV2 (0.8 ppm). Images were acquired with a saturation scheme of B_1_=2 μT, bandwidth=150 Hz, duty cycle=60% that generated minimal CEST contrast. (B) Images of AAV2 phantom (5.26×10^8^ vg/μL) at the same frequency offsets but with an optimized saturation scheme of B_1_=5 μT, bandwidth=50 Hz, duty cycle=85% that generated high CEST contrast. (C) Representative z-spectrum and 2-pool Lorentzian fitting of AAV2 from the low-contrast saturation scheme. (D) Representative z-spectrum and 2-pool Lorentzian fitting of AAV2 from the high-contrast saturation scheme with visible peak around 0.8 ppm. (E) Corresponding Lorentzian difference plot showing little to no CEST contrast for AAV2 around 0.8 ppm for low-contrast saturation scheme. (F) Corresponding Lorenzian difference plot displaying 11.85% CEST contrast around 0.8 ppm for high-contrast saturation scheme.

**Table 1.** Optimized CEST-MRI Saturation Parameters. CEST saturation schemes at 7 T tested that exhibit high levels of CEST contrast (>9%) in AAV2 phantom (5.26×10^8^ vg/μL).

| **B<sub>1</sub> (μT)** | **Bandwidth (Hz)** | **Duty Cycle (%)** | **Contrast (%)** |
| --- | --- | --- | --- |
| 3 | 50 | 0.7 | 9.42 |
| 3 | 50 | 0.85 | 9.96 |
| 4 | 75 | 0.7 | 10.44 |
| 4 | 75 | 0.85 | 10.63 |
| 4 | 50 | 0.7 | 11.48 |
| 4 | 50 | 0.85 | 11.35 |
| 5 | 50 | 0.7 | 11.28 |
| 5 | 50 | 0.85 | 11.82 |

Evaluation of CEST contrast generated by AAV2 in the presence of biological background signals yielded lower, but still detectable levels of contrast as expected. A phantom containing multiple AAV2 samples at different concentrations (5.26×10^3^ vg/μL-5.26×10^8^ vg/μL) suspended in cell lysate solution were imaged using the same saturation schemes that generated comparatively higher levels of CEST contrast as previously mentioned (Figure 7A). Subsequent analysis of CEST acquisitions yielded mean CEST values of 1.26±1.24%, 1.58±1.27%, 2.01 ±1.60%, and 3.39±1.93% across saturation schemes for AAV2 capsid concentrations of 5.26×10^3^ vg/μL, 5.26×10^5^ vg/μL, 5.26×10^7^ vg/μL, and 5.26×10^8^ vg/μL, respectively. Compared to the CEST-NMR results of AAV2 alone, detection of CEST contrast from AAV2 in biological media at 7 T still demonstrated a significant, albeit slightly weaker positive correlation between concentration and CEST contrast (R^2^=0.94, P=0.03). At the highest AAV2 concentration tested, optimized saturation schemes were able to generate CEST contrast levels of up to 5.85% (Figure 7B-F).

**Figure 7.**
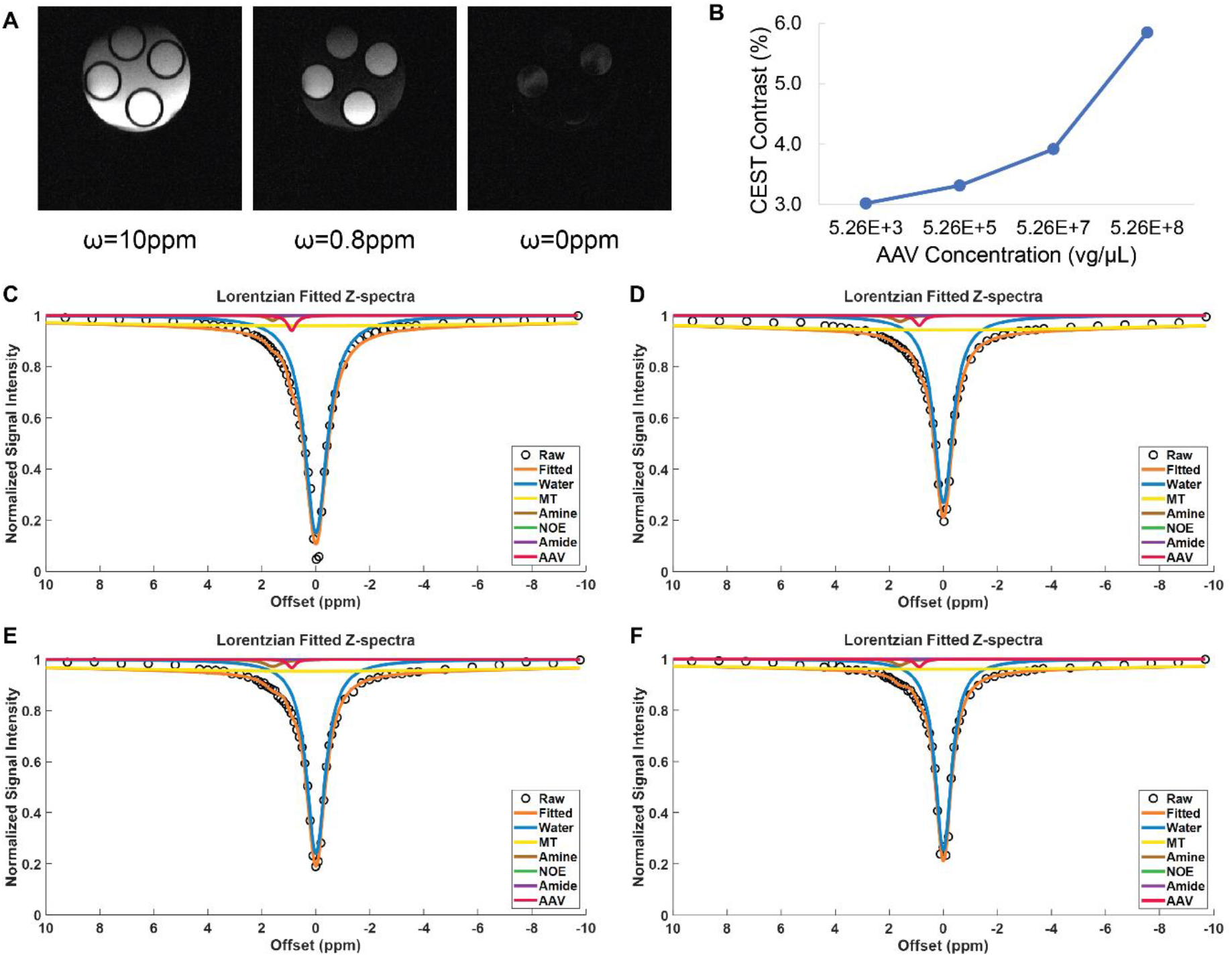
Z-spectra and Lorentzian Fitting of AAV2 in Presence of Background Signal. (A) Images of multi-concentration AAV2 phantom and cell lysate phantom at multiple frequency offsets that correspond to the highest acquired offset (10 ppm) and the resonant frequencies of water (0 ppm) and AAV2 (0.8 ppm). Concentrations included in the phantom were 5.26×10^8^ vg/μL (top), 5.26×10^7^ vg/μL (right), 5.26×10^5^ vg/μL (bottom), and 5.26×10^3^ vg/μL (left). (B) Resulting contrast across AAV2 concentrations tested using an optimized saturation scheme of B_1_=5 μT, bandwidth=50 Hz, duty cycle=70%. Z-spectrum and 6-pool Lorentzian fitting for AAV2 phantom at concentrations of 5.26×10^8^ vg/μL (C), 5.26×10^7^ vg/μL (D), 5.26×10^5^ vg/μL, and 5.26×10^3^ vg/μL.

## 4. Discussion

In this study we determined the resonance frequency of AAV2 and probed its corresponding levels of CEST contrast under several varying conditions, namely saturation power, pH, density, and biological transduction stage. AAV2 has a resonance frequency around 0.6-0.8 ppm and demonstrates robust contrast that scales up with saturation power until it reaches a plateau. CEST contrast of AAV2 viral capsids displays a significant exponential correlation with viral capsid density and a linear correlation with pH across saturation powers. Detection of AAV2 internalization across timepoints showed varying trends in both CEST contrast and viral titer/cell count but exhibited a strong positive correlation between viral titer/cell count and corresponding CEST contrast. Experiments conducted to explore CEST contrast of other AAV serotypes revealed slightly reduced but comparable levels of contrast around the same resonant frequency of AAV2. Following calculation of AAV2 exchange rates to be around 1300 Hz, subsequent optimization of a CEST saturation scheme for AAV2 at 7 T indicated multiple sets of saturation parameters that yield high levels of CEST contrast across saturation powers of 3-5 μT, bandwidths of 50-75 Hz, and duty cycles of 70-85%. Evaluation of AAV2 CEST contrast in the presence of biological background signal resulted in an expected reduction in observed signal that still correlated with concentration. Thorough investigation of CEST contrast generated by AAV2 viral capsids reveals promising results that bolster the potential of AAV2 as an endogenous CEST contrast agent.

Previous studies have explored various methods of assessing gene editing and the outcomes of gene therapy. Traditionally, verification of successful gene editing is performed via biopsy of the target tissue and sequencing for subsequent determination of gene editing efficiency^26,27^. Other methods for validation of gene editing utilize an imaging platform for a less invasive approach. PET and SPECT employ reporter genes to help visualize the spatial distribution and amplitude of gene expression following administration of gene therapy treatment. Previous studies have explored the use of PET in conjunction with the herpes simplex virus type 1-thymidine kinase (HSV1-TK) reporter for oncolytic virotherapy and cancer gene therapy^28–30^. By phosphorylating non-toxic prodrugs in infected cells to transform them into toxic forms that trigger apoptosis, HSV1-TK acts as a suicide gene that can also be used to image transgene expression^31^. In comparison, the human sodium iodide symporter (hNIS) reporter gene used in PET and SPECT studies operates by expressing the sodium iodide symporter for mediating active transport of iodide (I^-^), including radioisotope forms, thereby serving as both a reporter and therapeutic gene^32,33^. The hNIS gene has been used in multiple studies for monitoring tumor formation, inducing therapeutic effect, and evaluating gene expression patterns in animal models of heart failure and varying types of cancer^34–38^. However, the advantages of employing these reporter genes are to some extent offset by the use of ionizing radiation in both PET and SPECT imaging modalities, which can potentially lead to malignancies^39^.

Evaluation of the results of gene therapy has previously been performed with other imaging modalities as well, such as MRS and MRI. Facilitating the probing of metabolic changes associated with transgene activity, MRS has been applied to gene therapy studies to assess metabolite changes in tumors following siRNA-based gene therapy^40^ and in the brain following AAV vector-mediated gene delivery for targeting neurodegenerative diseases^41,42^. MRI allows monitoring of structural changes associated with transgene activity and has been used in previous studies for guiding delivery of an AAV gene therapy dosage as well as tracking subsequent gene expression in Parkinson’s disease, Alzheimer’s disease, and GM1-gangliosidosis^43–45^. A drawback of both MRS and MRI is how these imaging modalities primarily assess outcomes that occur following transgene expression without providing critical information on intermediate stages in the process of gene therapy. Furthermore, both magnetic resonance methods only probe one category of effects associated with gene therapy and thus often require the use of additional testing to fully verify the efficacy of treatment.

CEST-MRI was first applied to LRP, a nonmetallic, biodegradable reporter with an abundance of amide protons on the lysine that exchange with surrounding water^20^. Through frequency-selective saturation at +3.76 ppm relative to water, signal from LRP can hence be detected as a reduction in the water signal. LRP has since been explored in the settings of oncolytic virotherapy^31^ and rat gliomas^32^. In another study, an AAV9 vector encoding for LRP was delivered into mice hearts via direct injection and demonstrated significantly higher CEST contrast compared to empty vectors^27^.

Since the AAV capsid itself has demonstrated capacity for generating CEST contrast, it can potentially be used for noninvasive monitoring across the gene therapy process, including delivery, transduction, and gene expression in vivo, without the need for an additional tag or the use of ionizing radiation. Although we originally expected the source of CEST contrast from AAV to be from amide protons on lysine residues located on the capsid surface, the discovery of the resonant frequency associated with AAV to be approximately 0.6-0.8 ppm instead of around 3.5-3.7 ppm points to an alternative proton pool generating contrast. Based on previous studies investigating the structure of the AAV capsid, we believe the CEST signal from AAV can be attributed to hydroxyl protons from serine and threonine residues on the capsid surface^46–48^. The shift in resonant frequency in the range of 0.6-0.8 ppm is likely from background signal components in the endosomes and cell lysate that create a slightly different chemical environment compared to AAV simply in solution. Furthermore, the reduction in CEST contrast when moving from AAV samples in solution to AAV samples prepared in biological media is from dilution with other pools whose signals also contribute to corresponding Z-spectra. Analysis of CEST contrast from AAV2 may be subject to interference from neighboring proton pools, such as amine around 2 ppm, which could have potentially affected proper detection of signal from the viral capsid during the Lorentzian fitting process. However, the lower CEST contrast values in the presence of background signals are comparable to levels of contrast observed in other studies examining endogenous CEST effects^49–53^.

Several limitations to this study are worth noting. First, a limited number of AAV2 capsid concentrations were used for testing resulting CEST contrast at both the NMR and 7 T scanners. Given the large increase in CEST contrast generated between 5.26×10^7^ vg/μL and 5.26×10^8^ vg/μL, it may be useful to evaluate contrast at intermediate concentrations or past the highest concentration used in this study to examine to what extent CEST contrast continues to increase with increasing concentration. Second, our findings were all conducted using in vitro experiments and were not verified in vivo. Despite the inclusion of results from isolated endosomes and from phantoms with biological media, it would be useful to evaluate changes in CEST contrast of AAV in an animal model for further verification of its potential for gene therapy monitoring.

## 5. Conclusions

The primary findings of this study are that AAV2 viral capsids generates robust CEST contrast in vitro across a variety of chemical environments, concentrations, and saturation schemes. Significant correlations were observed for AAV2 CEST contrast when adjusting pH, concentration, and viral titer per cell count and should be taken into consideration for further investigation of contrast characteristics. Additional experiments to explore the effectiveness of AAV2 viral capsids as an in vivo endogenous CEST contrast agent for gene therapy tracking is needed.

## 6. Acknowledgements

This study was supported by the following funding sources: NIH 1R01HL28592, Silvian Foundation Award, NIH UH2EB028908, AHA 19TPA34850040, DGE 1752814.

---

**Supplemental Figure 1.**
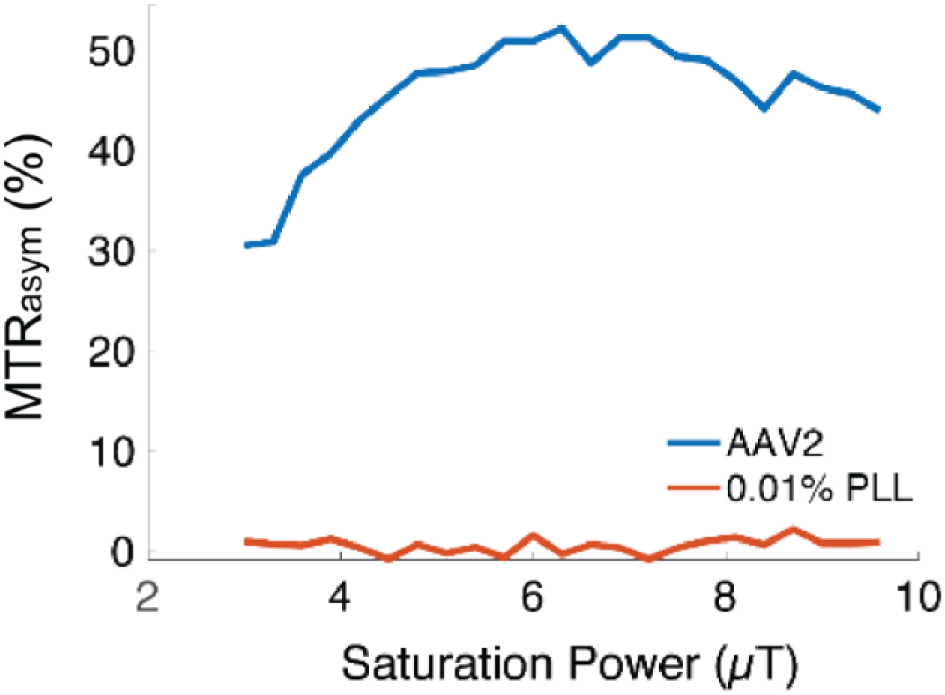
CEST Contrast Across Saturation Powers. Resulting CEST contrast of AAV2 (5.26×10^8^ vg/μL) and 0.01% PLL as a control across saturation powers ranging from 3 μT to 9.9 μT.

**Supplemental Figure 2.**
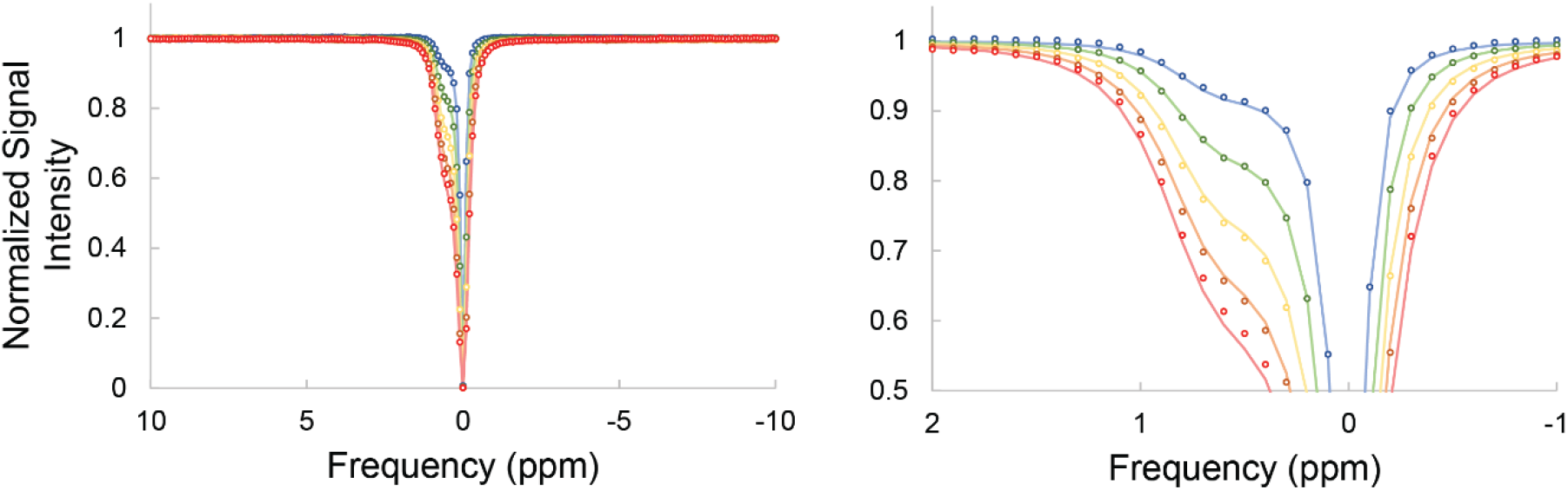
Bloch-McConnell Fitting for Exchange Rate Determination. Example of multi-B_1_ Bloch-McConnell equation fitting of z-spectra for determination of exchange rate. Plots shown are for pH 6.

